# Automated assembly of high-density carbon fiber electrode arrays for single unit electrophysiological recordings

**DOI:** 10.1101/2022.08.17.504264

**Authors:** Tianshu Dong, Lei Chen, Paras R Patel, Julianna M Richie, Cynthia A Chestek, Albert J Shih

## Abstract

**Objective:** Carbon fiber (CF) is good for chronic neural recording due to the small diameter (7 *µ*m), high Young’s modulus, and low electrical resistance, but most high-density carbon fiber (HDCF) arrays are manually assembled with labor-intensive procedures and limited by the accuracy and repeatability of the operator handling. A machine to automate the assembly is desired.

**Approach:** The HDCF array assembly machine contains: 1) a roller-based CF extruder, 2) a motion system with three linear and one rotary stages, 3) an imaging system with two digital microscope cameras, and 4) a laser cutter. The roller-based extruder automatically feeds single CF as raw material. The motion system aligns the CF with the array backend then places it. The imaging system observes the relative position between the CF and the backend. The laser cutter cuts off the CF. Two image processing algorithms are implemented to align the CF with the support shanks and circuit connection pads.

**Main results:** The machine was capable of precisely handling 6.8 *μ*m carbon fiber electrodes. Each electrode was placed into a 12 *μ*m wide trenches in a silicon support shank. Two HDCF arrays with 16 CF electrodes populated on 3 mm shanks (with 80 *μ*m pitch) were fully assembled. Impedance measurements were found to be in good agreement with manual assembled arrays. One HDCF array was implanted in the motor cortex in an anesthetized rat and was able to detect single unit activity.

**Significance:** This machine can eliminate the manual labor-intensive handling, alignment and placement of single carbon fiber during assembly, providing a proof-of-concepts towards fully automated HDCF array assembly and batch production.

## 1. Introduction

Electrophysiological recordings contribute to both the study of neurophysiological mechanisms and the optimal construction of future brain-machine interfaces [1–4]. Single neuronal firing generates internal voltages of 80 to 100 mV on a millisecond timescale. But external spikes can only be distinguished within an approximately 150 *μ*m radius [5]. Implantation of microelectrode arrays (MEAs) in the brain places recording sites close to neurons, providing high spatial and temporal resolutions for the electrophysiological recording of single neuron activity [6,7]. Thin diameter microwire-based MEAs are commonly used for single unit recordings. The thin microwire electrodes made of biocompatible platinum, tungsten, carbon, or stainless steel materials have both high Young’s modulus and high flexibility. While high Young’s modulus provides the rigidity for implantation, the microwires’ high flexibility mitigates the inconformity in stiffness of the brain and the electrode, making microwires suitable for chronic recording over several months [1–3,8–14]. In comparison, high rigidity silicon-based MEAs may be more likely to trigger an immune response generating glial scars followed by possible signal loss over time [15,16].

Among microwire materials used for electrophysiological recordings, carbon fiber (CF) has the smallest diameter of 5– 10 *μ*m [17], minimizing the immune responses. CF also has a low electrical resistivity, which facilitates the detection of low voltage signals without noise contamination [18]. These characteristics have made CF electrodes (CFEs) ideal for chronic recording [19–21].

Two configurations of CFE arrays are bundle and linear array. The bundle CFE array was first developed by Guitchounts et al. using 16 CFEs threaded by hand through a 3D-printed plastic block [13]. The CFE bundle was dipped in a water bath with surplus fiber section exposed in the air, and trimmed along the water surface by torch burning [19]. Based on the same technique, Lee et al. constructed a bundle of 4 CFEs and added a biodegradable silk fibroin support to facilitate the deeper penetration into the brain [22]. The bundle CFE array has a high local recording density but sacrifices the spatial coverage and the miniature size advantage of the CFE.

The linear CFE array could overcome the limitations of bundle CFE by using individual CFEs aligned in a 2D configuration with high spatial density. Research has been conducted to assemble high-density carbon fiber (HDCF) arrays both manually by hand and automatically by a machine. For manual fabrication, Patel et al. [1,23] constructed the first linear HDCF array of 16 CFEs with a 150 μm pitch. The array was constructed on a printed circuit board (PCB) with 16 gold traces. Silver epoxy was dispensed on the gold traces and the CF was placed in the epoxy pool manually under a microscope. The silver epoxy was cured at 140°C, held CFEs in place, and established the electrical connection between the CFE and PCB. A further study was conducted to build an HDCF array of 16 CFEs with 132 *μ*m pitch on a flexible polyimide PCB and a 532 nm laser was applied for CFE tip preparation instead of manual scissor cutting [24,25]. Huan et al. utilized a silicon support structure for the 16 CFE array with 80 μm pitch to achieve up to 9 mm insertion depth [20]. Individual CFs were cut to length and manually placed along the shanks in the silicon support structure using forceps. This manual assembly of CFE array is labor-intensive, time-consuming, and limited by the accuracy and repeatability of the operator.

For automatic fabrication of HDCF arrays, Massey et al. [26] proposed an automated micro-assembly method to build a 32-channel CFE array with 32 μm pitch. Two silicon-based microfabricated substrates, namely device and alignment substrates, were precisely aligned and stacked with micro-positioners, forming 32 holes filled with conductive silver ink and epoxy. CF was extruded from a roller-based extruder. An image detection algorithm was used to align and insert each CF into corresponding hole. CFs inserted in holes were aligned vertically to the substrates by raising the assistive substrate and then fixed inside holes by silver ink and epoxy curing. An array with a small pitch distance and high assembly accuracy was achieved after removing the assistive substrate. This machine demonstrated the feasibility of using image processing techniques for alignment between centers of CF circular bottom surfaces and device substrate holes. However, it required the usage of multiple silicon microfabricated substrates and micro-positioners for CF parallel alignment. Also, after automatic insertion of CF into each substrate hole, a tedious manual process was needed to cut off the CF with fine surgical micro-scissors. To overcome these drawbacks, the HDCF assembly machine developed in this study further advanced the image-based assembly concept and developed automatic contact-free CF cutting approach. By using a two-camera imaging system, the developed assembly system had the ability to detect and align individual CF in four degrees of freedom, including three-dimensional position of the CF tip and CF orientation for parallelism in a linear array. CF movement could thus be precisely guided solely based on image processing results. A laser-based CF cut off method was also integrated into the system for the first time, eliminating the manual mechanical cutting step.

In the present study, an HDCF array assembly machine was developed to automatically construct HDCF arrays with 80 *μ*m pitch. The precise placement of 6.8 *μ*m diameter CFEs along trenches (12 *μ*m in width) in silicon supporting shanks was performed under image guidance and correspondingly developed image processing algorithms. This machine could enable automatic alignment and placement of a single CF onto the HDCF array PCB with a specific length extended from the end of the silicon support. Subsystem tests on algorithms performance were conducted. Two HDCF arrays were assembled. The assembled HDCF arrays were evaluated by impedance test and in-vivo recording from rat motor cortex.

## 2. Methods

### 2.1 HDCF array

As shown in figure 1A, the targeted HDCF array for assembly automation was composed of a connector (A79040-001 by Omnetics, Minneapolis, MN), a custom PCB, a silicon support structure (SSS), and 16 CFEs. The connector had standardized pins for back-end electrical connection to pre-amplifiers. SSS was fabricated with the six-step cleanroom procedure introduced in [20]. The SSS, as shown in figure 1B, provided physical and electrical connections between CFEs and custom PCB and had four components: 1) silicon base and shanks, 2) bond pads to PCB, 3) gold traces, and 4) gold pads, as well as two indented features: 1) silver epoxy wells beneath each gold pad and 2) trenches along the middle line of each shank. The length and density of CFE in HDCF array were pre-determined by the length and pitch of the shanks. In this study, the shanks were 3 mm long and the pitch was 80 μm. CFEs extended about 3.5 mm from the silver epoxy wells and formed an array with 80 μm spacing.

**Figure 1.**
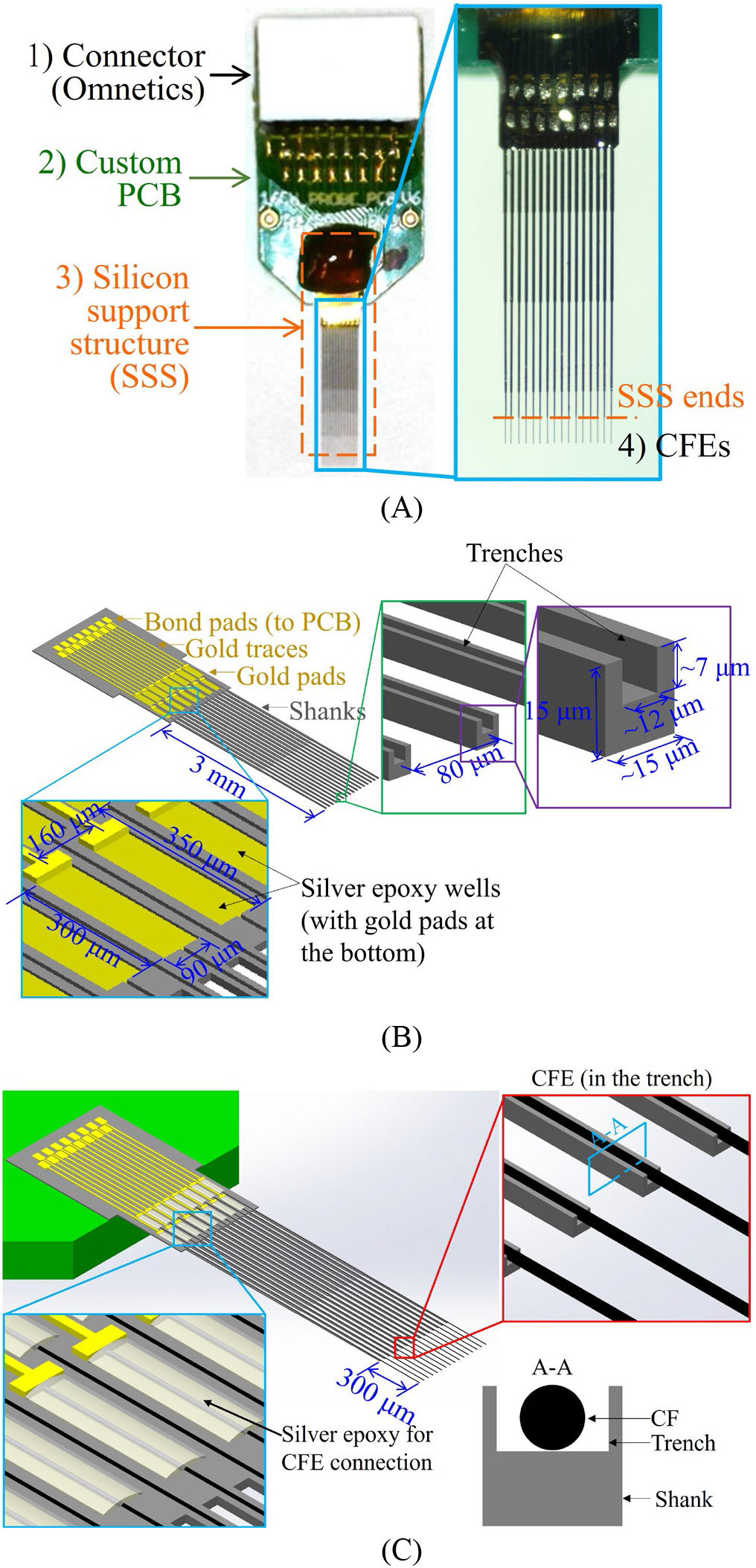
(**A**) HDCF array components: connector, custom PCB, SSS, and CFEs, (**B**) SSS components, features, and dimensions, and (**C**) CFEs started from the silver epoxy wells and overhangs from the trenches.

During assembly of HDCF array, the CFEs were placed on the top of gold pads, laid in the trenches and supported by the shanks in the SSS. Each CFE overhangs by about 0.3 mm beyond the end of the shank trenches, as shown in figures 1A and 1C, and had non-insulated recording sites at the free end tips. The CFEs and shanks of SSS were implanted together, providing a minimally invasive solution for the deep insertion of CFE into recording sites. The rigidity of SSS ensured the successful insertion of CFEs. The minimal invasiveness was a result of the small 15 *μ*m by 15 *μ*m cross-sectional area of the CFE and the shank (figures 1B and 1C) [20].

### 2.2 HDCF array assembly

Seven steps were taken to assemble an HDCF array.

Step 1 Connection of connector, PCB, and SSS. The connector, PCB, and SSS were connected physically and electrically. The method used here was elaborated in [20].

Step 2 CF loading. A CF section (about 6 cm in length) was loaded onto the roller-based extruder, elaborated in Section 2.3.1.

Step 3 CFE population. 16 CFEs were placed in silver epoxy wells and trenches with the HDCF array assembly machine. Details of the machine and the process will be illustrated in Sections 2.3 to 2.5.

Step 4 CFE electrical connection. Silver epoxy (H20E; Epoxy Technology, Billerica, MA) was deposited in silver epoxy wells by a desktop nanolithography system (NLP 2000, Advanced Creative Solutions Technology, Carlsbad, CA) and then cured at 140 °C for 20 minutes. CFEs were electrically connected to the SSS and PCB.

Step 5 CFE physical reinforcement. Epoxy (NOA 61, Norland Products, Cranbury, NJ) was applied on the gold pad area and along the shanks and cured under UV light to physically reinforce the connection between CFEs and the SSS.

Step 6 Device insulation. The HDCF array was conformal coated with approximately 800 nm of Parylene C (PDS 2035, Specialty Coating Systems, Indianapolis, IN) for insulation.

Step 7 CFE functionalization. CFEs were laser cut to 0.3 mm overhanging length using a 532nm Nd:YAG pulsed laser (LCS-1, New Wave Research, Fremont, CA; 5 mJ/pulse, 5 ns pulse duration), plasma treated using a Glen 1000P Plasma Cleaner (pressure 200 mT, power 300 W, time 120 s, oxygen flow rate 60 sccm, and argon flow rate 7 sccm), and coated with poly(3,4-ethylenedioxythiophene):sodium p-toluenesulfonate (PEDOT:pTS) as previously done in [1,14]. The short, 0.3 mm, overhanging (unsupported) length of CFEs avoids buckling during implantation.

### 2.3 HDCF array assembly machine

The HDCF array assembly machine developed in this study is shown in figure 2. It consisted of four subsystems: 1) a roller-based extruder for CF feeding, 2) a motion system for PCB translation and CF rotation, 3) an imaging system for motion trajectory planning and process monitoring, and 4) a laser cutter for CF cutoff. The Subsystems 1-4 are described in the following Sections 2.3.1 to 2.3.4. The controller and software platform applied to coordinate the subsystems are presented in Section 2.3.5. The target workpiece for the machine was the PCB with SSS, which was attached to the motion system by a 3D-printed fixture.

**Figure 2.**
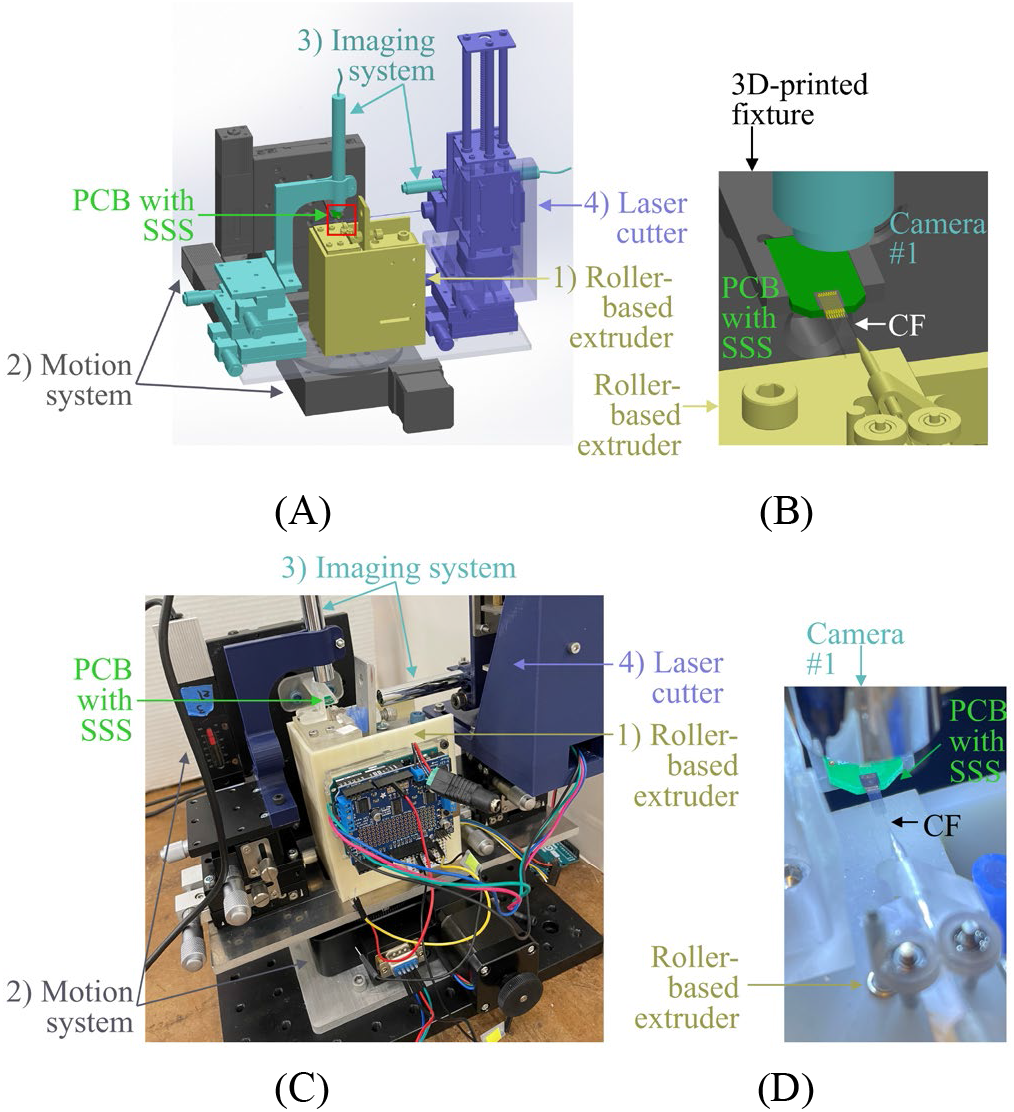
HDCF array assembly system and its four subsystems: (**A**) 3D overview of the 1) roller-based extruder, 2) motion system, 3) imaging system, and 4) laser cutter, together with the PCB with SSS as the starting workpiece, **(B)** close-up view of the red-boxed area in figure 2A including the PCB with SSS, camera (in the imaging system), and CF (from the roller-based extruder); pictures of **(C)** the physical machine and (**D**) close-up view of the roller-based extruder, the PCB with SSS, Camera #1, and the CF.

#### 2.3.1 Subsystem 1: Roller-based extruder

The roller-based extruder realized automatic feeding of single CF for the HDCF array assembly system. As shown in figure 3, this subsystem consisted of: 1) an adjustable base, 2) two feeding rollers, 3) a direct current (DC) drive motor, and 4) two Parylene C coated tapered glass capillary tubes. Two main features distinguished this CF feeding extruder from the open-source automated high-density microwire neural recording array assembly system [21]: 1) adjustable roller compression preload for feeding, 2) coated capillary tubes to prevent CF sticking due to electrostatic charging.

**Figure 3.**
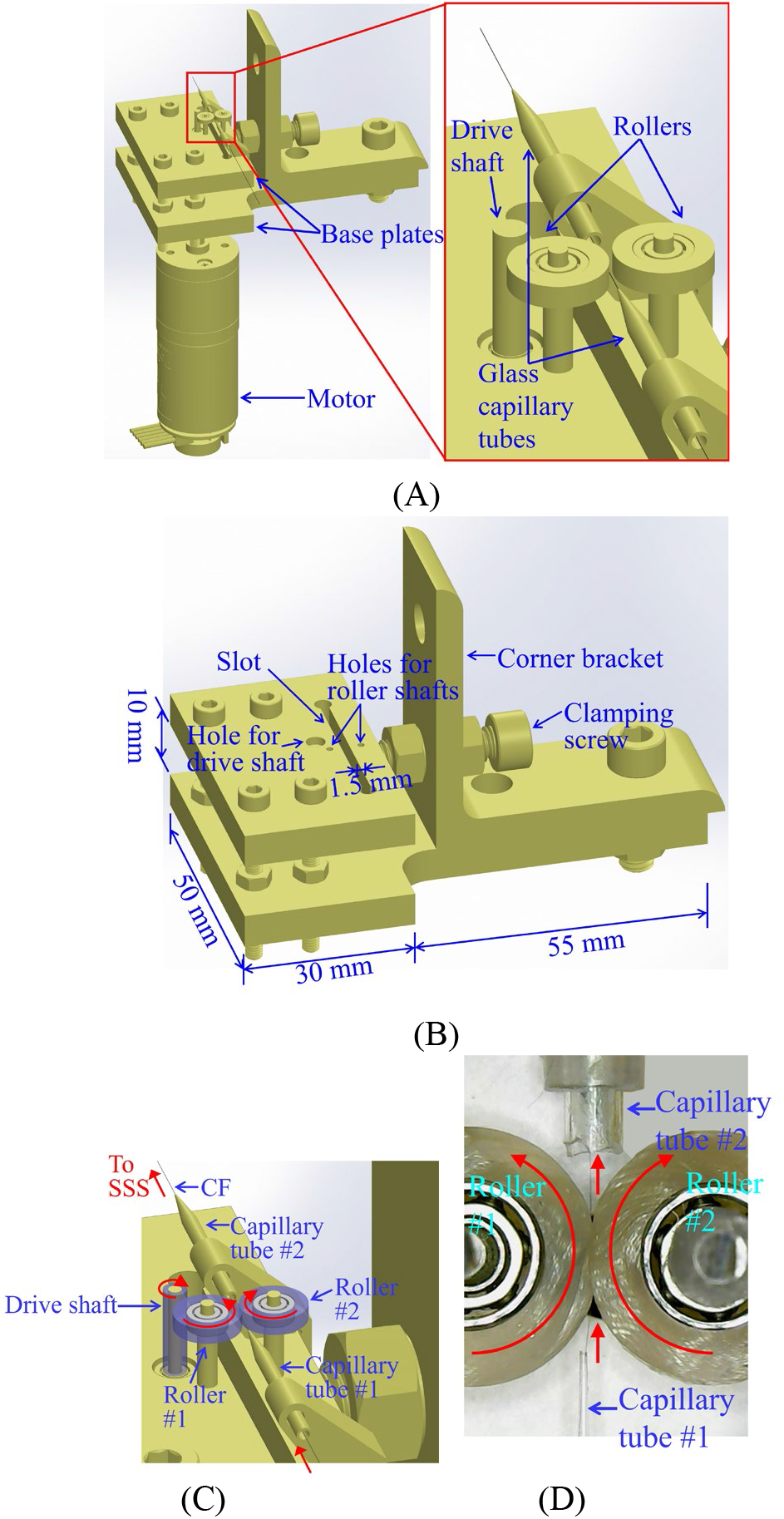
Roller-based extruder: (**A**) overview of the adjustable base, the rollers, the motor, and the glass capillary tubes, (**B**) the adjustable base, (**C**) rollers and shaft with soft silicone coating (colored in transparent azure) and CF extrusion mechanism (motions are marked in red arrows), and (**D**) a photo of CF being advanced from capillary tube #1 to capillary tube #2 by the friction from the rollers.

The adjustable base (figure 3B) contained two base plates 10 mm apart, to provide structural support for the rollers and the motor shaft. Both plates were 3D-printed (Form 2, Formlabs Inc., Somerville, MA) with transparent stereolithography resin (RS-F2-GPCL-04, Formlabs, Somerville, MA). The top plate had a 1.5 mm wide slot parallel and close to the edge, forming a flexural beam structure. Two rollers (5.4 mm in diameter) were located on the opposite side of this slot. The surfaces of the two feeding rollers and the motor drive shaft were wrapped by a layer of soft silicone (durometer 50A hardness) to increase the frictional force along the feed direction and move CF at the roller contact surface. By adjusting the clamping screw, the roller shaft distance and thus roller compressive preload could be precisely controlled.

Two tapered glass capillary tubes were used to provide guidance for the CF along the feeding direction. The two tubes were set along the perpendicular bisector of the two roller’s centers with their tips pointing towards the feeding direction. CF would be advanced through capillary tube #1, between two rollers, then through capillary tube #2 towards the PCB, as shown in figures 3C and D.

Our study showed that for the capillary tube without coating, five attempts on CF loading/extrusion were made, and none of them succeeded due to CF adhering, likely by electrostatic force, to the inner wall of the glass capillary tube. For the capillary tube with Parylene C coating, extrusion tests on four CFs (15 extrusions per CF) were performed with 100% success (n = 60). This indicated that the Parylene C coating on the glass capillary tube reduced the adhesion force and the static friction. Thus, the inner walls of glass capillary tubes were coated with approximately 800 nm of Parylene C through chemical vapor deposition to prevent the CF-tube adhesion due to electrostatic charge.

To load the CF into the extruder for automatic feeding, a long section (about 6 cm in length) of CF was manually threaded into capillary tube #1 until the CF tip touched the contact region of the two rollers. The motor (Model 2285, Pololu, Las Vagas, NV) was turned on and its drive shaft (1.8 mm in diameter) in contact with roller #1 drives both rollers under a synchronized rotation through friction. The CF was advanced to the outlet of capillary tube #2 and the extruder was ready for use to feed CFEs onto SSS of the HDCF array.

To test the accuracy of the extruder, 11 CF extrusion tests were carried out. The desired length of extrusion was set as 4 mm. The length of extruded CF was evaluated based on images taken by the imaging subsystem (Section 2.3.3) before and after extrusion. The results showed good accuracy with extrusion length of 4.009 ± 0.073 mm (n = 11, mean ± SD).

#### 2.3.2 Subsystem 2: Motion system

The motion system had one rotary stage (HT03RA100, Jiangyun Technology, Beijing, China) and three linear stages (100cri-R, Siskiyou, Grants Pass, OR), as shown in figure 4A. The rotary stage rotated the roller-based extruder to get the extruded CF parallel with the shanks in the SSS. The center of rotation was selected as the tip of the capillary tube #2 of the roller-based extruder. The rotary stage also rotated the two digital cameras in the imaging system and the laser cutter together to keep the CF-camera-laser relative position and alignment. The *x* and *y* linear stages then translated the PCB with the information from the imaging system (Section 2.3.3) and image processing algorithms (Section 2.4) to align a CF with the specific trench in the shank of the SSS along the *x*- and *y*-directions. Finally, the *z* linear stage elevated the PCB to land the CF tip on the gold pad in the SSS and place the CF within the trench in the shank of the SSS (to be elaborated in Section 2.5).

**Figure 4.**
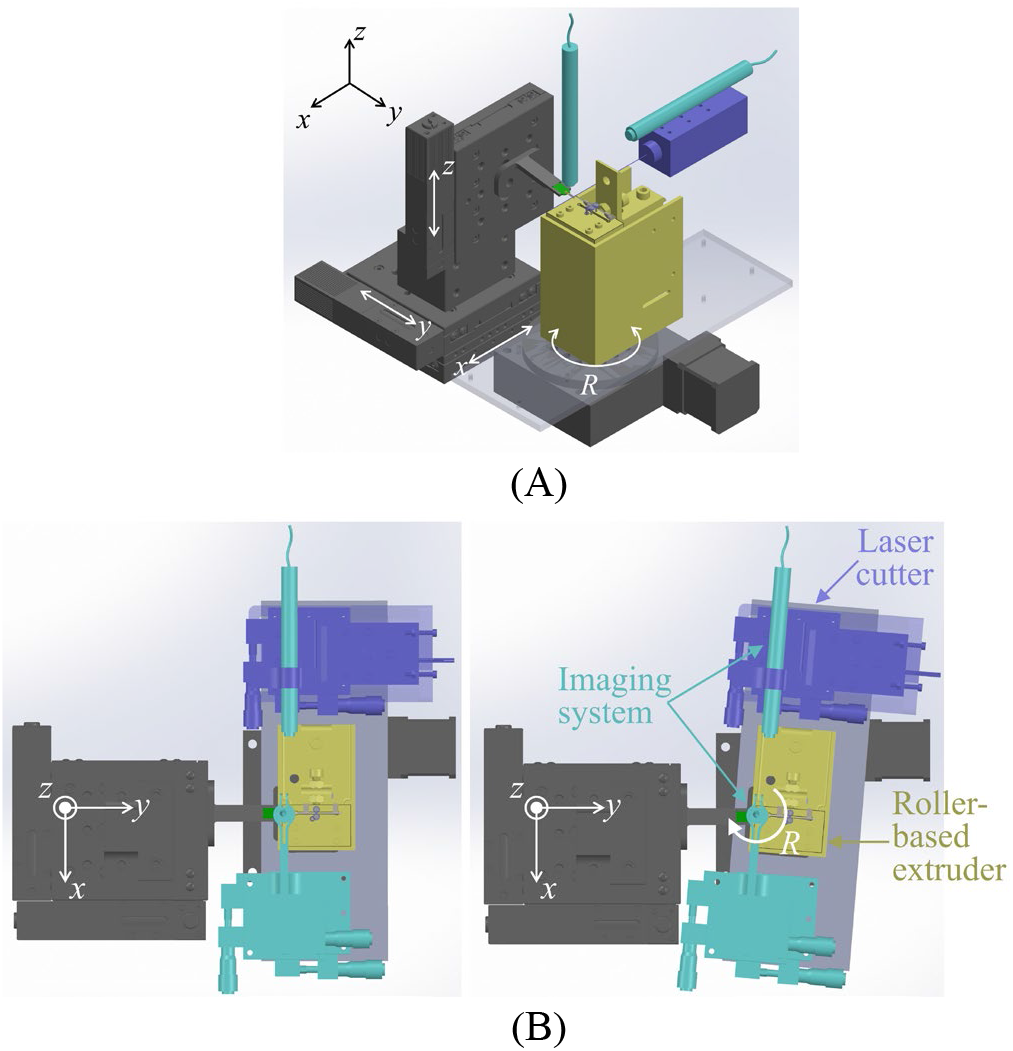
(**A**) Motion system: *x, y*, and *z* linear stages for PCB translation in *x*-, *y*- and *z*-directions and the rotary stage R for CF rotation and (**B**) the rotary stage rotates the roller-based extruder, the imaging system, and the laser cutter together (top view).

#### 2.3.3 Subsystem 3: Imaging system

Two digital microscope cameras (OT-HD, Opti-TekScope, Chandler, AZ), marked as cameras #1 and #2, are shown in figure 5A. Camera #1 was mounted on the rotating stage platform through a manual 3-axis linear stage (LD60-LM, ToAuto, Shenzhen, China), and pointed along the negative *z*-direction. The focus of camera #1 was adjusted to the extruded CF tip. Camera #1 provided an image of shanks and CF orientation in the *xy* plane. This image was the input for the image processing algorithms to find the CF tip’s relative position to the gold pads (figure 5B). Camera #2 was located on the side of the PCB and pointed along the positive *x*-direction with its focus on the shanks. Camera #2 provided the image of the shanks and CF orientation in the *yz* plane. This image was used to find the height of the PCB and determine the contact of CF with the SSS (figure 5C).

**Figure 5.**
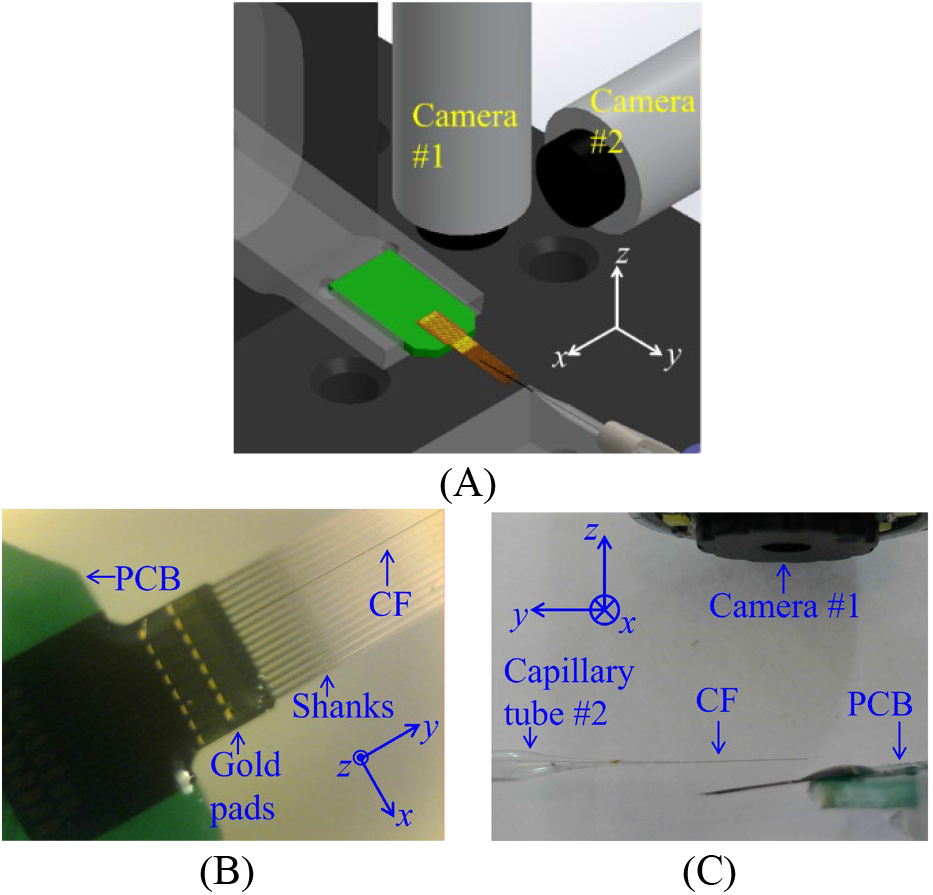
Imaging system: (**A**) camera #1 and camera #2, (**B**) top view of assembly area from camera #1, and (**C**) side view of assembly area from camera #2.

#### 2.3.4 Subsystem 4: Laser cutter

The laser cutter subsystem, as shown in figure 6A, consisted of 1) a blue laser head with 500 mW output power and 445 nm (± 5 nm) wavelength (D-B500F, Lian Dian Chuang Technology, Shenzhen, China), 2) a motorized linear stage (T8-Z150, Huizhou Bachin Electronic Technology, Huizhou, China), and 3) a manual 3-axis linear stage (LD60-LM, ToAuto, Shenzhen, China). The position of the laser head in *y*-direction was adjusted to align with the Capillary #2 tip with the manual linear stage prior to assembly.

**Figure 6.**
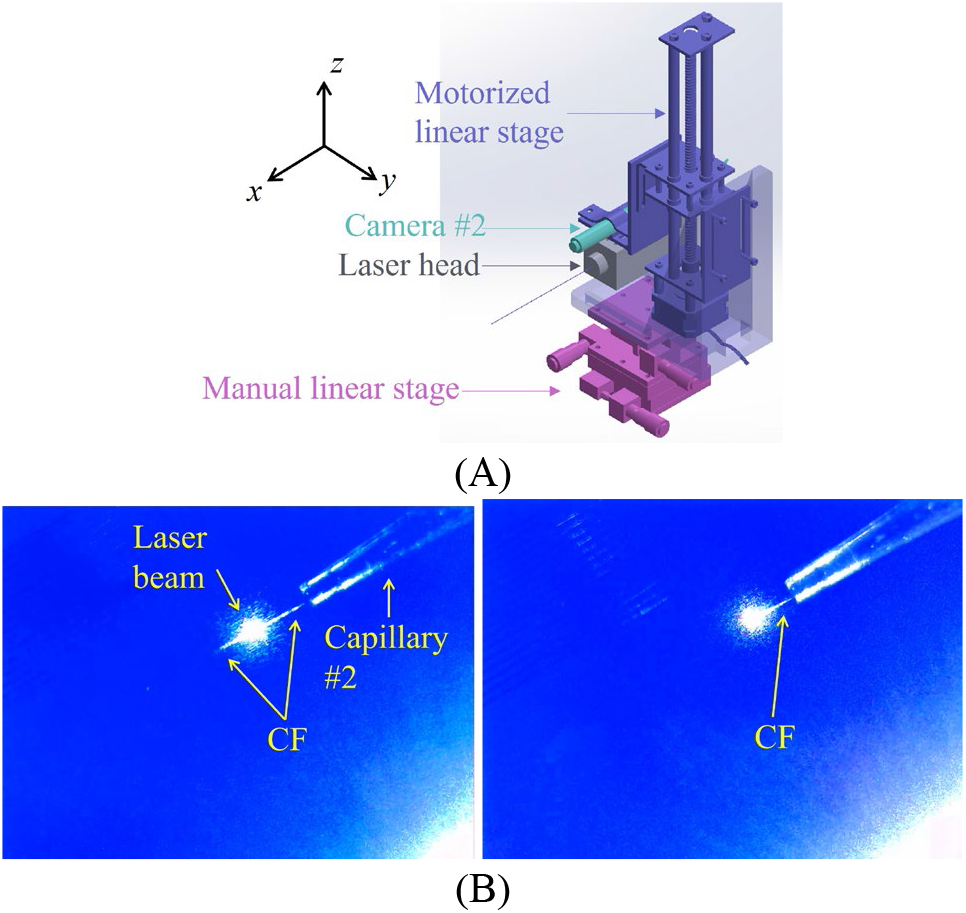
(**A**) Laser cutter components: laser head, motorized linear stage, and manual linear stage and (**B**) laser beam cutting off the CF (top view).

To cut off the CF from the extruder, the laser head was powered on and moved by the motorized linear stage in the positive *z*-direction until the laser beam hit the CF, as shown in figure 6B. The CF could be cut off in less than five seconds with the 12 DCV input voltage of the focused laser beam. After cutting, the laser head was moved down to its original position to allow the camera #2 for imaging.

#### 2.3.5 Controller and software platform

MATLAB (R2020b, Mathworks, Natick, MA) was used as the master software platform for computer control and synchronization of all subsystems in the HDCF array assembly machine. The roller-based extruder was controlled using an Arduino Uno microcontroller. The controller of three linear stages was NI PCI-7344 with LabVIEW VIs (National Instruments, Austin, TX). The controller of the rotating stage and the motorized linear stage for the laser cutter was the Arduino Uno microcontroller with a DRV8825 stepper motor driver.

### 2.4 Algorithms for CFE population

Two algorithms were developed to achieve the automatic CFE population. To be placed precisely into the trench and silver epoxy well, the CFE needed to be aligned parallel with the shank first, then its tip should contact the center of a gold pad edge. Algorithm I measured the orientation of CF and shanks. Algorithm II calculated the distance between the CF tip and the desired edge center on gold pads where the tip would be placed. Both algorithms used the digital images taken by Camera #1 from the top of the CF and the PCB as input. Details for Algorithms I and II are presented in the next two sections.

#### 2.4.1 Algorithm I: CF-shank orientation alignment

The goal of Algorithm I was to measure the orientation of the shank and CF for their angular alignment. Two images used as input for Algorithm I were the image of the gold pad area and part of the shanks from camera #1 for measuring shank orientation *β*_1_ and the image of the CF for measuring the CF orientation *β*_2_. The CF and shanks were orientated diagonally as shown in figure 7 to view a longer section of both. The output *β* = *β*_2_ – *β*_1_ was the difference between the two measured angles. The CF would be rotated by an angle *β* using the rotary stage in the motion system for the CF-shank alignment.

**Figure 7.**
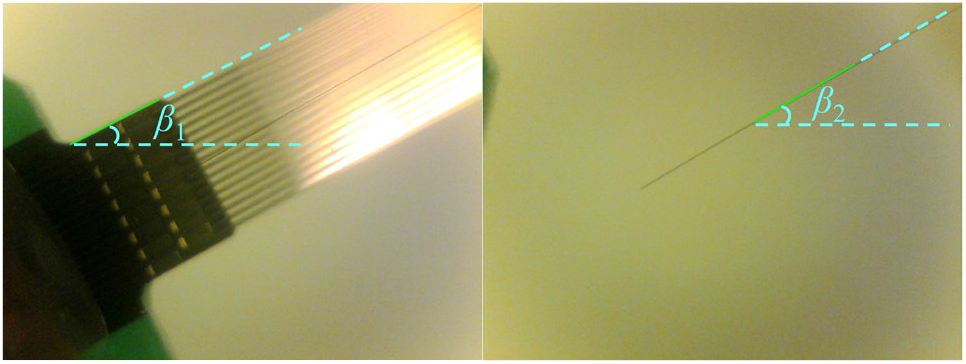
Measurement of shank orientation *β*_1_ and CF orientation *β*_2_.

#### 2.4.2 Algorithm II: CF tip – gold pad edge center alignment

Algorithm II calculated the distance between the desired gold pad edge center position and the CF tip and decomposed it into *x*- and *y*-displacements in two orthogonal directions for the translational motion of the PCB. The input for Algorithm II was the digital image of the gold pad area and CF tip location. The CF tip pixel location was tagged manually as *O* (see figure 8) and used as the origin for the Cartesian coordinate for the PCB translational motion. *O* remained still in the view during CF rotation and PCB translation because the camera was fixed with respect to the CF. The output variables were the distances Δ*x* and Δ*y*, as shown in figure 8, between the CF tip and the target (highlighted in yellow) in *x*- and *y*-directions, respectively. For each of the sixteen rectangular gold pads, the target was the middle point of the shorter edge away from the shanks. The CF tip would be aligned with the target and placed in the trench of SSS. In this way, when silver epoxy was deposited on the gold pads after CFE placements, a long section of CFE would be in contact with silver particles in the epoxy to ensure proper electrical connection.

**Figure 8.**
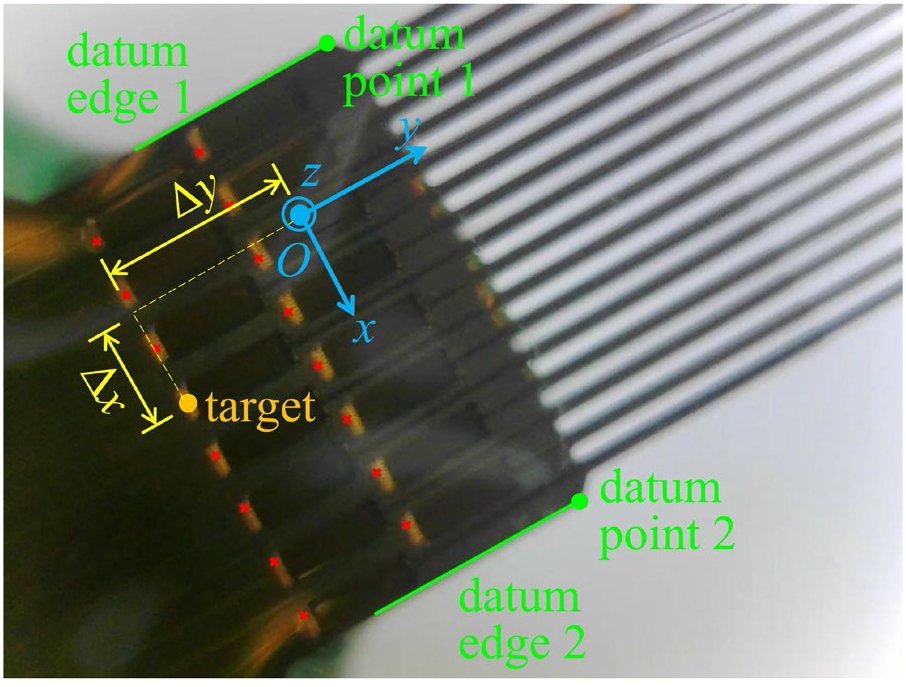
Algorithm II: inputs (blue), outputs (yellow), and datum points and edges (green).

To calculate Δ*x* and Δ*y* from the image, two mappings had to be constructed. These were 1) between the line orientation in the image and the directions for PCB translation and 2) between the pixel distance in the image and the distance in PCB translational motions. To build such connections, two datum edges, the upper and lower bounds of the gold pad area, and two datum points, the corners of the gold pad area, were extracted using MATLAB. The orientation of the datum edges in the image was mapped to the *y*-direction for PCB translation and the pixel distance was mapped to the width of the gold pad area. Positions of centers of the gold pad’s edges could be calculated based on the datum edges, points, and design sketch of the silicon support structure. The pixel distance of the CF tip and target gold pad edge’s center was first decoupled into *x*- and *y*-directions, then converted into distance Δ*x* and Δ*y*.

### 2.5 CFE population

To facilitate CFs sliding into the trenches, a drop of deionized water was put on and spread across the SSS with a tapered glass capillary tube. As shown in figure 9, around the epoxy which secures the SSS to the PCB (refer to Section 2.2 Step 1), a liquid pool was formed. The deionized water pool was thick at the gold pad area and shallow around the shank. During the CFE population, deionized water needed to be refilled whenever it totally evaporated. At the completion of the CFE population, deionized water had evaporated and no solid residue was left on the SSS.

**Figure 9.**
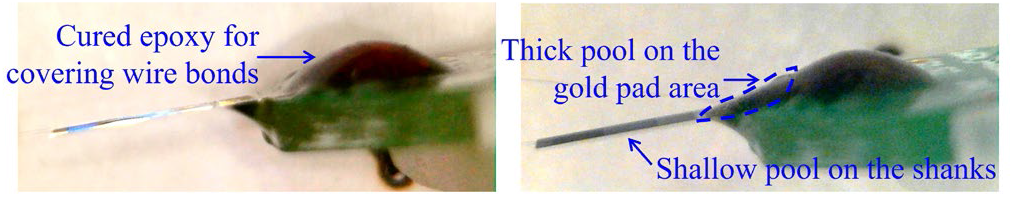
Photos from camera #2 before and after the deionized water application around the epoxy covering the wire bonds between the SSS and PCB.

CFEs were populated on the SSS shank-by-shank by repeating an extrusion-placement-cut off process. First, a 4 mm long CF was fed from the tip of capillary tube #2. The CF tip is located above the gold pad area and inside the viewing area of Camera #1. Second, CF was aligned parallel to the shank based on Algorithm I, and its tip was moved to the target position based on Algorithm II, which was executed 3 times iteratively. Third, the CF tip was landed on the gold pad and the rest of the CF was pulled into the trench by surface tension, as shown in figure 10. Lastly, the CF was cut off with the laser cutter after verifying that the full length of the shank was occupied by CF in camera #2’s view. The extrusion-placement-cutoff process is repeated 16 times for the CFE population in 16 gold pads on the SSS.

**Figure 10.**
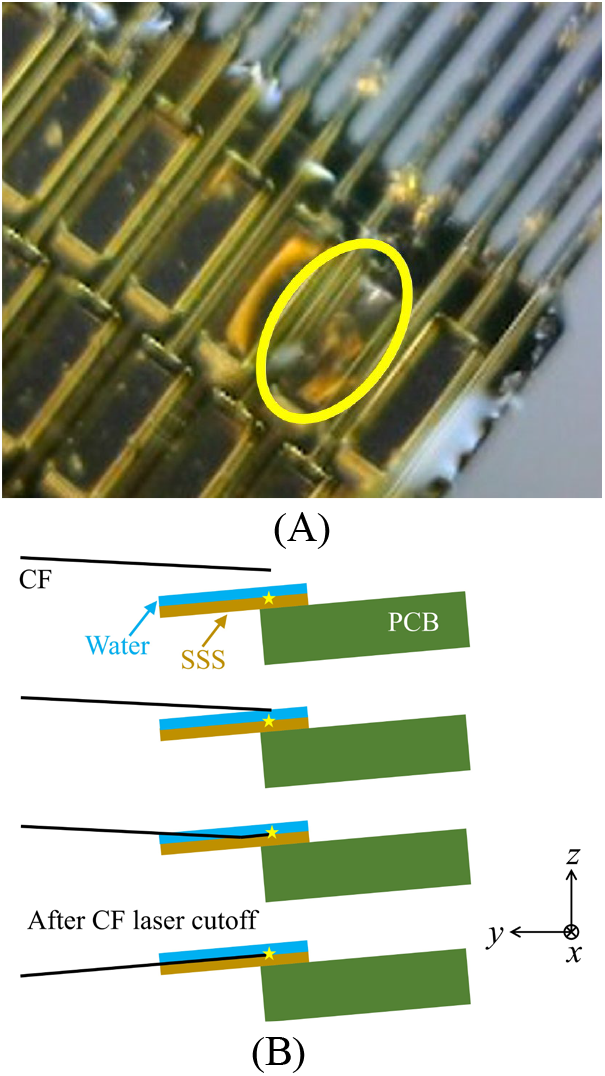
CF landing: (**A**) CF tip touching the gold pad and (**B**) CF pulled into the trenches on the shanks by the surface tension of the water.

### 2.6 Electrochemical impedance spectroscopy (EIS) and electrodeposition

To characterize the arrays’ impedances, measurements were taken with a PGSTAT12 Autolab potentiostat (Metreohm/Eco Chemie, Utrecht, Netherlands) which was controlled using the vendor supplied NOVA software. For each measurement session, the device was submerged by approximately 1 mm in 1x phosphate buffered saline (BP3994, Fisher, Waltham, MA). The reference was provided by an Ag|AgCl electrode (RE-5B, BASi, West Lafayette, MA) and the counter by a stainless steel rod. Measurements were applied from 10 Hz to 31 kHz with a 10 µV_RMS_ signal. Data was analysed using custom MATLAB scripts. The deposition of PEDOT:pTS is described in [24].

### 2.7 Electrophysiology recordings and neural data analysis

To test the functionality of the devices *in vivo*, a male Long Evans rat was first anesthetized using a combination of ketamine (90 mg/kg) and xylazine (5 mg/kg) via intraperitoneal (IP) injection. Carprofen (5 mg/kg) was given once, subcutaneously. Subsequent boosters of ketamine (30 mg/kg) were also given IP. Once down the rat’s head was shaved, then cleaned using alternating wipes of 70% ethanol and betadine. Next a midline incision was made and the skin was pulled back. A bone screw in the posterior portion of the skull was screwed in and served as the ground/reference connection. Next a craniotomy was made over the motor cortex, the dura resected, and the probe inserted. All procedures and monitoring complied with the University of Michigan’s Institutional Animal Care & Use Committee. Recordings, described in detail [14], were taken with a system from Tucker-Davis Technologies. Electrophysiology data analysis was conducted with Offline Sorter (Plexon, Dallas, TX) with the full methodology described in [24].

## 3. Results

### 3.1 CF deviation angle reduction with Algorithm I application

We first evaluated how well the CF could be aligned parallel to the silicon supports. After aligning CFs using the apparatus, 15 sets of CF deviation angles (angle between the CF and the shank in the top view, denoted by *θ*_xy_) before and after Algorithm I execution were measured and compared. The *θ*_xy_ was measured three times with the Fiji image processing package [27] in ImageJ (v.1.52r). The average value of *θ*_xy_ before Algorithm I application was 1.16 ± 0.94 degrees (mean ± SD) and was reduced to 0.54 ± 0.51 degrees after the adjustment. The alignment in the orientation between CF and the shank is improved with the Algorithm I application.

### 3.2 CF tip positioning accuracy using Algorithm II

We next evaluated how precise the machine could place the CF tip to the target point on the gold pad. CF tip positioning accuracy after applying Algorithm II was evaluated by measuring the distance (error) between the CF tip *O* and the target in *x*- and *y*-directions (denoted as *ε*_x1_ and *ε*_y1_) after moving the PCB by Δ*x* and Δ*y* calculated by Algorithm II. A total number of 48 measurements were performed. The results of *ε*_x1_ vs. Δ*x* and *ε*_y1_ vs. Δ*y* are summarized in figure 11. Due to image distortion further from the center, the error in tip position in the *x*-direction *ε*_x1_ increased as the absolute value of Δ*x* increased. The *ε*_x_ larger than 40 μm would end up as a failed assembly because CF tip would be outside of the gold pad. The gold pad had a large dimension (300 μm) in the *y*-direction, the *ε*_y1_ ranging from 0 to 100 μm was tolerable.

**Figure 11.**
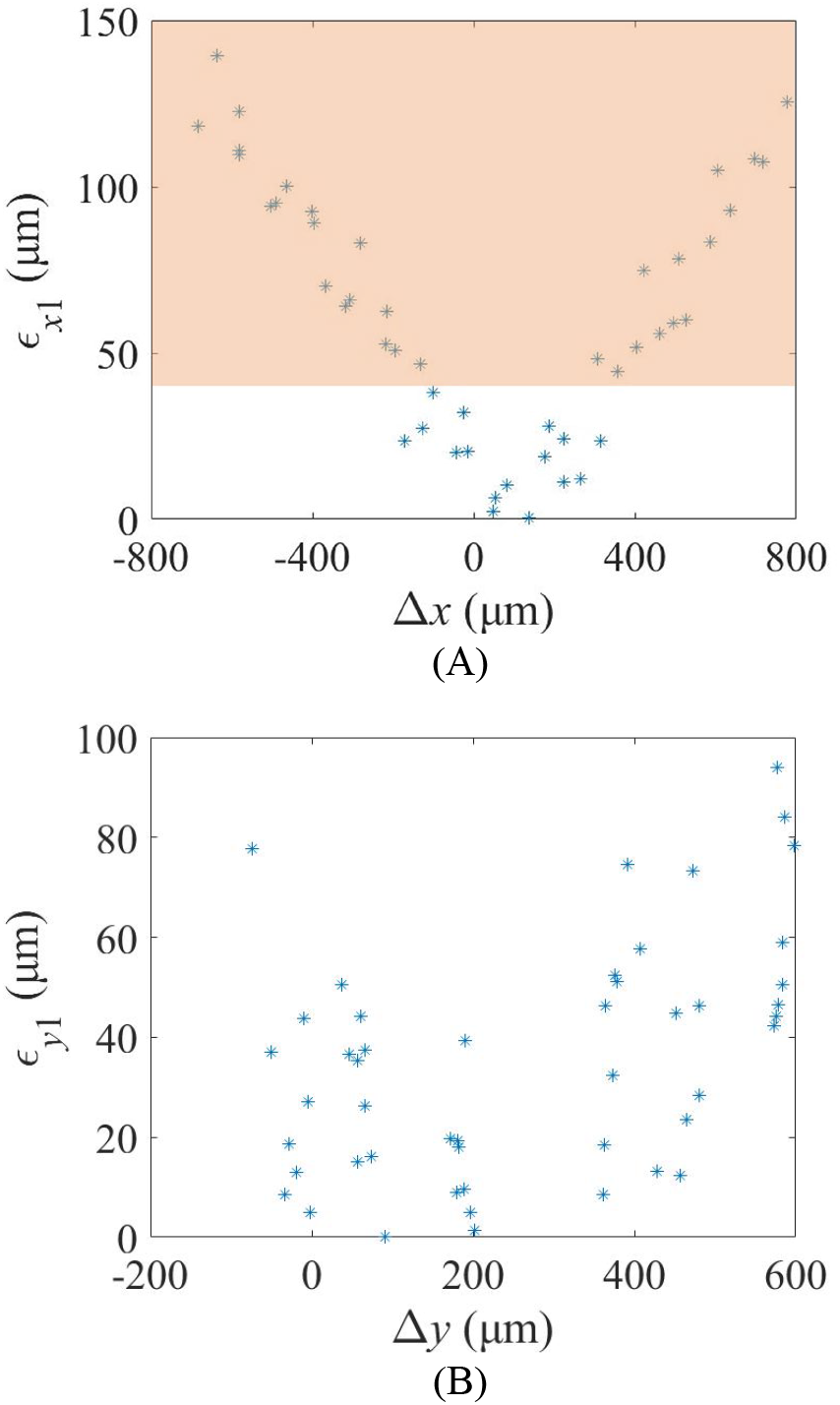
CF tip positioning accuracy in (**A**) *x*- and (**B**) *y*-directions after first iteration of Algorithm II. The failure area is shaded with *ε*_x1_ larger than 40 μm.

After the first iteration using Algorithm II, the CF tip moved to a position closer to the center of the target gold pad with less image distortion. The Algorithm II was applied again based on positions from the first iteration. Errors were measured and denoted as *ε*_x2_ and *ε*_y2_. The changes of *ε*_x1_ and *ε*_x2_ of the 48 measurements are summarized in figure 12A. For 45 out of the 48 cases, the error in the *x*-direction was reduced to less than 40 μm after the second iteration. For the three cases with *ε*_x2_ greater than 40 μm, a third iteration of Algorithm II was utilized and the CF tip was aligned within the gold pad with the error *ε*_x3_ all less than 30 μm. For HDCF machine operation, Algorithm II was operated 3 times.

**Figure 12.**
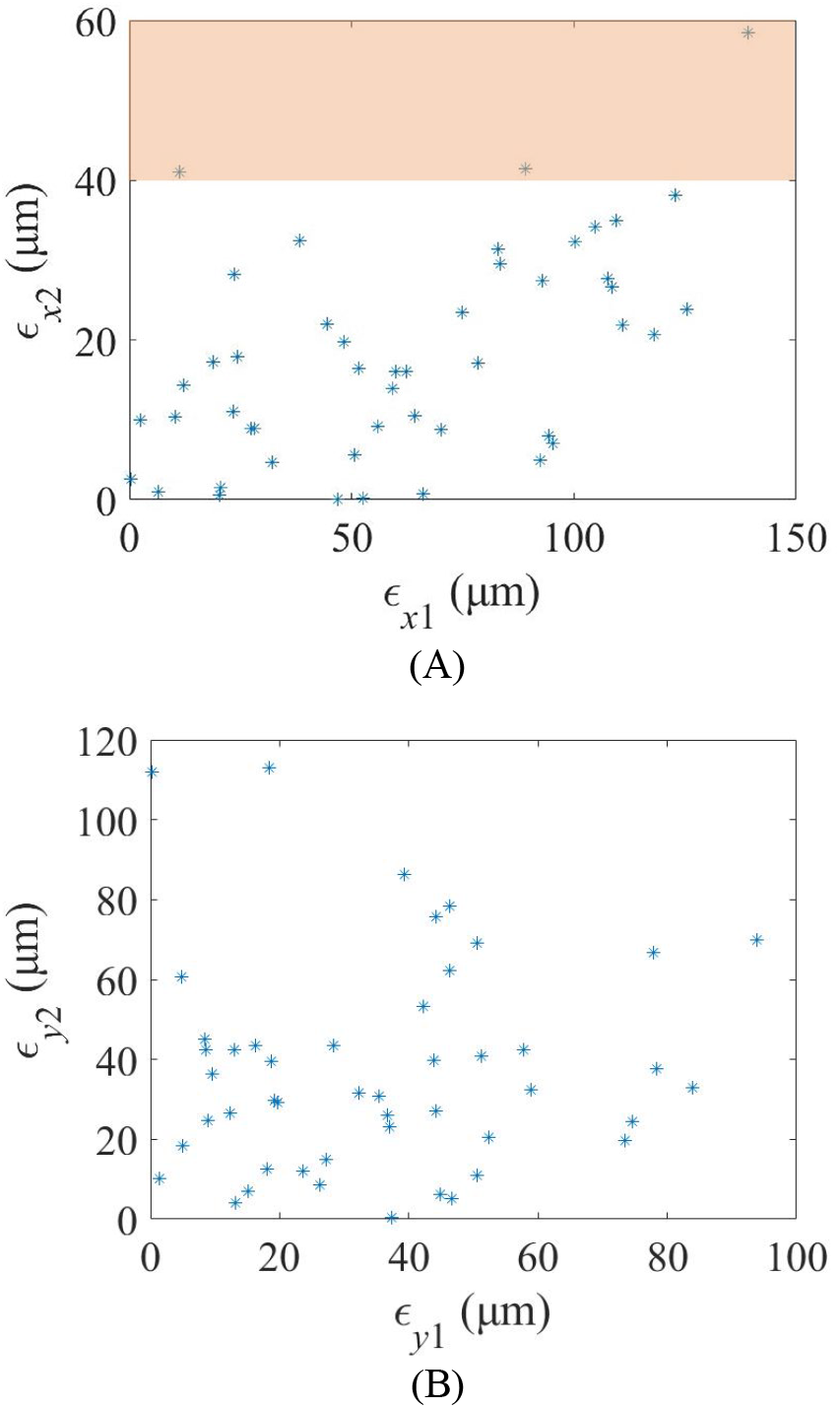
Error of CF tip position from the gold pad center after the second iteration of repeated Algorithm II: (**A**) *ε*_x2_ vs. *ε*_x1_ (the failure area is shaded with *ε*_x2_ larger than 40 μm) and **(B)** *ε*_y2_ vs. *ε*_y1_.

### 3.3 Successful population of arrays with 16 CFEs

Two HDCF arrays with 16 CFEs populated on 3 mm shanks (with 80 μm pitch) were fully assembled using this machine, as shown in figure 13. All 16 CFEs were placed accurately within the silver epoxy wells and trenches to assemble the HDCF arrays and demonstrate the feasibility and repeatability of the HDCF array assembly machine.

**Figure 13.**
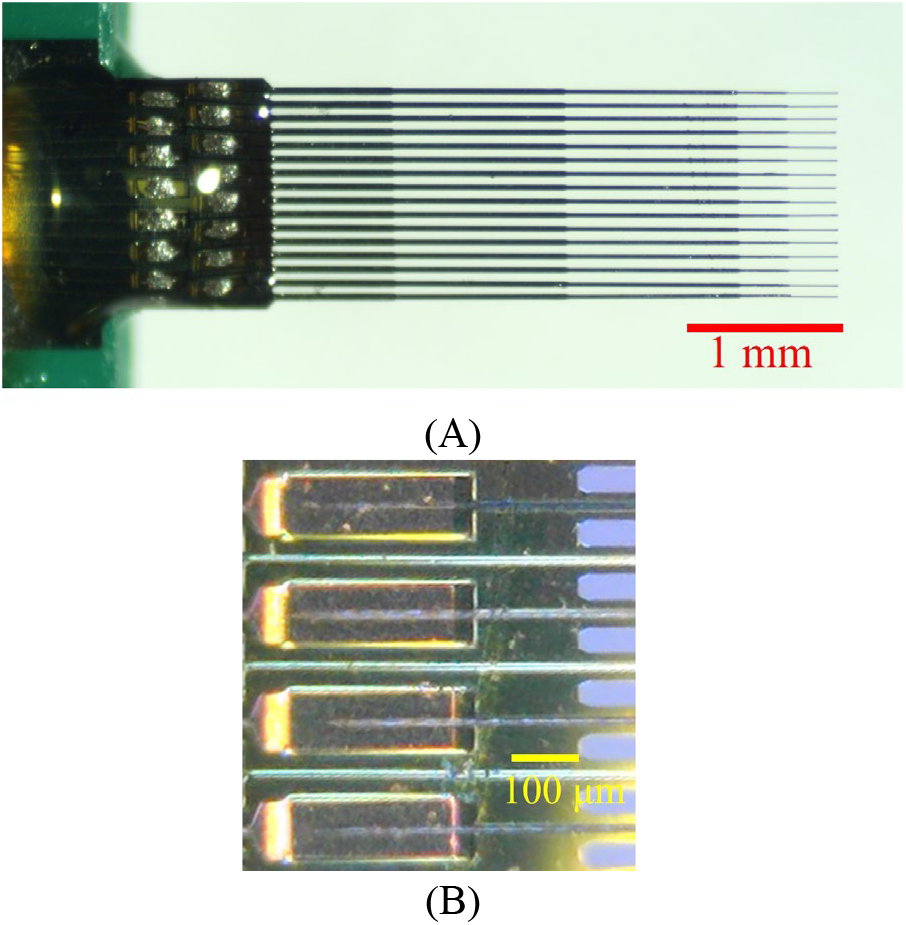
(**A**) Fully populated HDCF array and (**B**) the closed-up view of CFEs on the gold pads (before silver epoxy application).

### 3.4 EIS and in-vivo electrophysiology recording

After placement of CFs, additional steps were carried out to functionalize the arrays. After each major step listed below impedance measurements were taken. In brief, the silver epoxy was applied to the wells, heat cured, and the protruding CFs were cut by a scissor to 350 μm (n = 27 connected CFs, 164 ± 28 kΩ, 1 kHz impedance). Then the devices were coated with Parylene C, cut by laser to 300 μm overhanging length, exposed the tip [24], and treated by plasma (1682 ± 158 kΩ) [25]. Finally the newly exposed CF tips were coated with PEDOT:pTS via electrodeposition (18 ± 1 kΩ) [24]. The impedance measurements were found to be in good agreement with our past findings (figure 14) [24,25].

**Figure 14.**
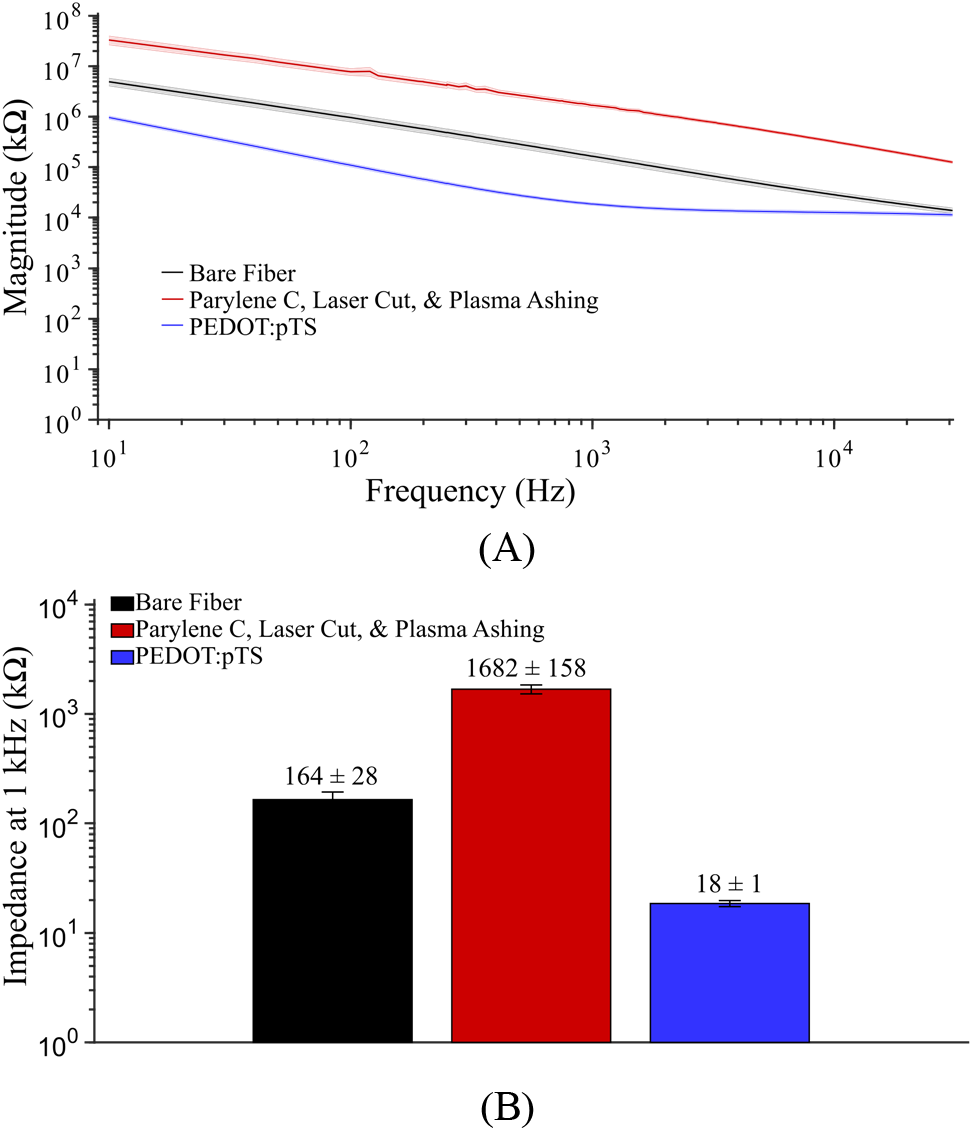
(**A**) Bode magnitude plots of the measured impedance after the three major functionalization steps (n = 27, mean ± SD). **(B)** One kilohertz impedance values (n = 27, mean ± SD) shown for the three major processing steps. As seen in previous results, coating with PEDOT:pTS greatly reduces the impedance of the electrodes.

To validate the fully assembled and functionalized HDCF array devices, one of them was implanted in layer V of the motor cortex in an anesthetized rat. Of the 16 channels present, 10 were able to detect single unit activity (figure 15). Across those 10 channels, 17 individual units were detected with a mean amplitude of 105 μV.

**Figure 15.**
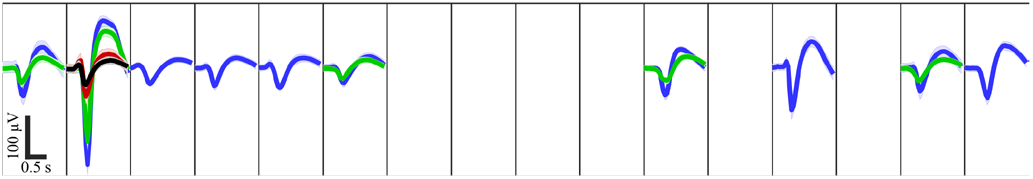
Spike panel of the detected unit activity from layer V motor cortex of an anesthetized rat.

## 4. Discussions

Here, we constructed an HDCF array assembly machine that precisely placed 6.8 μm CFEs with 80 μm pitch into 12 μm wide trenches, reducing the manual handling and microscope observation workload for the array assembly process. Our results here provide proof of concept for the possibility of automatic assembly CF arrays with minimal human operation and less specialized personnel.

The parameters in the elaborated algorithms may need manual tuning to reach an optimal detection. The integrated parameters were selected through trials and worked for over 80% of the detection trials. The failure might be a result of the fluctuating lighting conditions and was fixed by searching for two parameters following some strategy within the nearby range manually. This tuning process could be later automated according to the elaborated strategy. The parameters may also be selected by collecting the information of lighting condition through the background pixels and calculating the optimal value. The machine may work better in a controlled environment.

There are multiple implications of this work. First, like recent work using additive manufacturing [28], this HDCF array assembly machine demonstrates another alternative to traditional silicon array fabrication. While this work utilized CF, it could likely be extended to fabrication in other geometries and with other electrode materials not amenable to lithography such as tungsten. Second, this machine was relatively inexpensive and could be implemented in neuroscience labs that struggle with manual assembly of microwire arrays. Finally, the primary benefit of HDCF arrays compared to other softer subcellular electrodes is ease of surgical insertion, which simply requires a stereotaxic frame and micromanipulator. This paper suggests that a robotic approach at time of fabrication may obviate the need for a robot to perform the surgery itself.

## Acknowledgement

This work was financially supported by the National Institutes of Health National Institute of Neurological Disorders and Stroke (UF1NS107659). The authors would also like to thank Su Sung Kim who assisted in assembly tests.

## References

[1] Patel P R, Na K, Zhang H, Kozai T D Y, Kotov N A, Yoon E and Chestek C A 2015 Insertion of linear 8.4 μm diameter 16 channel carbon fiber electrode arrays for single unit recordings J. Neural Eng. 12

[2] Xie K, Fox G E, Liu J and Tsien J Z 2016 512-Channel and 13-Region Simultaneous Recordings Coupled with Optogenetic Manipulation in Freely Behaving Mice Front. Syst. Neurosci. 10

[3] Schwarz D A, Lebedev M A, Hanson T L, Dimitrov D F, Lehew G, Meloy J, Rajangam S, Subramanian V, Ifft P J, Li Z, Ramakrishnan A, Tate A, Zhuang K Z and Nicolelis M A L 2014 Chronic, wireless recordings of large-scale brain activity in freely moving rhesus monkeys Nat. Methods 11 670–6

[4] Mukamel R and Fried I 2012 Human Intracranial Recordings and Cognitive Neuroscience Annu. Rev. Psychol. 63 511–37

[5] Henze D A, Borhegyi Z, Csicsvari J, Mamiya A, Harris K D and Buzsáki G 2000 Intracellular Features Predicted by Extracellular Recordings in the Hippocampus In Vivo J. Neurophysiol. 84 390–400

[6] Thach W T and Bastian A J 2004 Role of the cerebellum in the control and adaptation of gait in health and disease Prog. Brain Res. 143 353–66

[7] Burle B, Spieser L, Roger C, Casini L, Hasbroucq T and Vidal F 2015 Spatial and temporal resolutions of EEG: Is it really black and white? A scalp current density view Int. J. Psychophysiol. 97 210–20

[8] Rose J D and Weishaar D J 1979 Tapered tungsten fine-wire microelectrode for chronic single unit recording Brain Res. Bull. 4 435–7

[9] Du Z J, Kolarcik C L, Kozai T D Y, Luebben S D, Sapp S A, Zheng X S, Nabity J A and Cui X T 2017 Ultrasoft microwire neural electrodes improve chronic tissue integration Acta Biomater. 53 46–58

[10] Williams J C, Rennaker R L and Kipke D R 1999 Long-term neural recording characteristics of wire microelectrode arrays implanted in cerebral cortex Brain Res. Protoc. 4 303–13

[11] Nicolelis M A L, Dimitrov D, Carmena J M, Crist R, Lehew G, Kralik J D and Wise S P 2003 Chronic, multisite, multielectrode recordings in macaque monkeys Proc. Natl. Acad. Sci. 100 11041–6

[12] Prasad A, Xue Q S, Sankar V, Nishida T, Shaw G, Streit W J and Sanchez J C 2012 Comprehensive characterization and failure modes of tungsten microwire arrays in chronic neural implants J. Neural Eng. 9 056015

[13] Guitchounts G, Markowitz J E, Liberti W A and Gardner T J 2013 A carbon-fiber electrode array for long-term neural recording J. Neural Eng. 10

[14] Patel P R, Zhang H, Robbins M T, Nofar J B, Marshall S P, Kobylarek M J, Kozai T D Y, Kotov N A and Chestek C A 2016 Chronic in vivo stability assessment of carbon fiber microelectrode arrays J. Neural Eng. 13 066002

[15] Polikov V S, Tresco P A and Reichert W M 2005 Response of brain tissue to chronically implanted neural electrodes J. Neurosci. Methods 148 1–18

[16] Karumbaiah L, Saxena T, Carlson D, Patil K, Patkar R, Gaupp E A, Betancur M, Stanley G B, Carin L and Bellamkonda R V. 2013 Relationship between intracortical electrode design and chronic recording function Biomaterials 34 8061–74

[17] MatWeb 2012 Solvay Thornel® T-650/35 3K Carbon Fiber, Polyacrylonitrile (PAN) Precursor

[18] Budai D 2010 Carbon fiber-based microelectrodes and microbiosensors Intelligent and Biosensors pp 269–88

[19] Guitchounts G and Cox D 2020 64-Channel Carbon Fiber Electrode Arrays for Chronic Electrophysiology Sci. Rep. 10 1–9

[20] Huan Y, Gill J P, Fritzinger J B, Patel P R, Richie J M, Della Valle E, Weiland J D, Chestek C A and Chiel H J 2021 Carbon fiber electrodes for intracellular recording and stimulation J. Neural Eng. 18 066033

[21] Massey T L, Lee J H, Ray M, Sathe N S, Liu X, Pister K S J and Maharbiz M M 2016 Open-source automated system for assembling a high-density microwire neural recording array 2016 Int. Conf. Manip. Autom. Robot. Small Scales, MARSS 2016

[22] Lee Y and Jun S B 2020 Carbon fiber based microeletrode with biodegradable silk for in vivo neural recording 2020 International Conference on Electronics, Information, and Communication, ICEIC 2020 (Institute of Electrical and Electronics Engineers Inc.)

[23] Patel P R 2015 Carbon FIiber Microelectrode Arrays for Neuroprosthetic and Neuroscience Applications (University of Michigan)

[24] Welle E J, Patel P R, Woods J E, Petrossians A, Della Valle E, Vega-Medina A, Richie J M, Cai D, Weiland J D and Chestek C A 2020 Ultra-small carbon fiber electrode recording site optimization and improved in vivo chronic recording yield J. Neural Eng. 17 026037

[25] Patel P R, Popov P, Caldwell C M, Welle E J, Egert D, Pettibone J R, Roossien D H, Becker J B, Berke J D, Chestek C A and Cai D 2020 High density carbon fiber 3arrays for chronic electrophysiology, fast scan cyclic voltammetry, and correlative anatomy J. Neural Eng. 17

[26] Massey T L, Santacruz S R, Hou J F, Pister K S J, Carmena J M and Maharbiz M M 2019 A high-density carbon fiber neural recording array technology J. Neural Eng. 16

[27] Schindelin J, Arganda-Carreras I, Frise E, Kaynig V, Longair M, Pietzsch T, Preibisch S, Rueden C, Saalfeld S, Schmid B, Tinevez J Y, White D J, Hartenstein V, Eliceiri K, Tomancak P and Cardona A 2012 Fiji: An open-source platform for biological-image analysis Nat. Methods 9 676–82

[28] Ali M A, Hu C, Yttri E A, Panat R, Ali M A, Hu C, Panat R and Yttri E A 2022 Recent Advances in 3D Printing of Biomedical Sensing Devices Adv. Funct. Mater. 32 2107671

